# The *677C>T* variant in *methylenetetrahydrofolate reductase* causes morphological and functional cerebrovascular deficits in mice

**DOI:** 10.1101/2021.12.16.472805

**Authors:** Alaina M. Reagan, Karen E. Christensen, Leah C. Graham, Amanda A. Bedwell, Kierra Eldridge, Rachael Speedy, Lucas L. Figueiredo, Scott C. Persohn, Teodoro Bottiglieri, Michael Sasner, Paul R. Territo, Rima Rozen, Gareth R. Howell

## Abstract

Vascular contributions to cognitive impairment and dementia (VCID) particularly Alzheimer’s disease and related dementias (ADRDs) are increasing; however, mechanisms driving cerebrovascular decline are poorly understood. Methylenetetrahydrofolate reductase (MTHFR) is a critical enzyme in the folate and methionine cycles. Variants in *MTHFR*, notably *677C>T*, are associated with dementias, but no mouse model existed to identify mechanisms by which *MTHFR^677C>T^* increases risk. Therefore, MODEL-AD created a novel knock-in (KI) strain carrying the *Mthfr^677C>T^* allele on the C57BL/6J background (*Mthfr^677C>T^*) to characterize morphology and function perturbed by the variant. Consistent with human clinical data, *Mthfr^677C>T^* mice have reduced enzyme activity in the liver and elevated plasma homocysteine levels. MTHFR enzyme activity as well as critical metabolites in the folate and methionine cycles are reduced in the *Mthfr^677C>T^* brain. Mice showed reduced tissue perfusion in numerous brain regions by PET/CT as well as significantly reduced vascular density and increased GFAP-expressing astrocytes in frontal cortex. Electron microscopy revealed cerebrovascular damage including endothelial and pericyte apoptosis, reduced luminal size, and increased astrocyte and microglial presence in the microenvironment. Collectively, these data suggest critical perturbations to cerebrovascular function in *Mthfr^677C>T^* mice supporting its use as a model for preclinical studies of VCID.

## Introduction

Vascular contributions to cognitive impairment and dementia (VCID) including Alzheimer’s disease (AD) are on the increase, yet the mechanisms defining vascular dysfunction are not well understood. AD is one of the most common forms of age-related neurodegenerative dementia^1^. Currently, over 50 million people globally are suffering from AD, and this number is expected to triple by 2050^2^. Multiple complex factors play a role in disease development, and it is now well-established that both genetic and environmental risk contribute to pathology^3, 4^. Historically, AD has primarily been characterized by the accumulation of beta amyloid plaques, neurofibrillary tangles of TAU and widespread neuronal loss^5^. Over the last 10 years, over 400 clinical trials for AD have failed, with the majority of them targeting amyloid and TAU accumulation^6^. Recent evidence suggests that the majority of cases of AD also present clinically with cerebrovascular complications that contribute to initiation and progression of the disease through diverse mechanisms including ischemia, vascular inflammation, blood brain barrier (BBB) dysfunction, small vessel disease and white matter hyperintensities – features shared with other dementias such as mixed dementia and vascular dementia (VaD)^7, 8^. To date, few clinical trials have attempted to improve cerebrovascular health to reduce risk or treat dementia, and currently there are a limited number of animal models that precisely manipulate cerebrovascular health to permit in-depth investigations, thus ultimately limiting our understanding of disease etiology.

Genome-wide association studies have linked the 677C>T polymorphism (often referred to as *C677T*) in the methylenetetrahydrofolate reductase (*MTHFR^677C>T^*) gene^9^ with both AD^10–12^ and vascular dementia^13^. Examining its contribution to AD is broadly relevant because 20-40% of the global population is predicted to be either heterozygous or homozygous for *MTHFR^677C>T^* ^14, 15^. Historically, null or highly deleterious mutations in *MTHFR* have been identified in patients with severe MTHFR deficiency due to an inborn error of metabolism resulting in homocystinuria^16^. Clinically, these *MTHFR*-deficient patients demonstrate vascular dysfunction, including thromboses^17^ and vascular lesions^18^. The *MTHFR^677C>T^* polymorphism is predicted to result in a less severe condition that has been linked to increased risk of many common peripheral vascular diseases in adulthood including stroke^19^ and atherosclerosis^20^. Additionally, clinical imaging studies have linked *MTHFR^677C>T^* to brain volume deficits in patients with mild cognitive decline^21^. Those who carry the risk variant consistently demonstrate reduced MTHFR enzyme activity^22^ and global hypomethylation^23, 24^ in leukocytes, and elevated plasma homocysteine^25–27^. Previously, MTHFR deficiency has been modeled using mice heterozygous and homozygous for a *Mthfr* null allele^28^, but the mechanisms by which the *MTHFR^677C>T^* variant specifically increases risk for VCID are not clear.

The *MTHFR* gene codes for a key regulatory enzyme in folate and homocysteine metabolism, and under physiological conditions generates the folate derivative (5-methyltetrahydrofolate) that is utilized for the conversion of homocysteine to methionine. Methionine is used to generate S-adenosylmethionine (SAM), a critical donor of methyl groups for numerous biological functions. Reduced MTHFR activity prevents homocysteine conversion, resulting in elevated homocysteine levels in the blood^29, 30^. This elevation has long been associated with vascular damage in cardiovascular disease, with evidence pointing to increased inflammation and endothelial cell dysfunction^31, 32^. Diets that induce hyperhomocysteinemia in mice also show VCID relevant phenotypes^33–35^. Diet modification has been used in clinical trials to lower blood homocysteine levels; although some studies reported a reduction in brain atrophy^36, 37^, trials were generally unsuccessful in modulating cognitive decline, suggesting a plasma homocysteine-independent mechanism of damage in the brain^38, 39^. Therefore, to better understand the role of the *MTHFR^677C>T^* variant in VCID, we created and characterized the *Mthfr^677C>T^* mouse strain that phenocopies decreased liver MTHFR enzyme activity and increased plasma homocysteine. In addition, *Mthfr^677C>T^* mice showed cerebrovascular deficits by PET/CT, immunohistochemistry and electron microscopy. These features suggest *Mthfr^677C>T^* alters cerebrovascular morphology and function, providing a new model in which to study mechanisms of VCID.

## Materials and Methods

All experiments were conducted in accordance with the ARRIVE guidelines.

### Mouse Strains

Animals were generated at The Jackson Laboratory (JAX) as part of the IU/JAX/Pitt/Sage MODEL-AD Center. Experiments were conducted in accordance with policies and procedures described in the Guide for the Care and Use of Laboratory Animals of the National Institutes of Health^40^ and were approved by the JAX Institutional Animal Care and Use Committee. All mice are congenic on the C57BL/6J (JAX# 000664) (B6) strain and bred and housed in a 12/12-hour light/dark cycle and fed LabDiet 5K52 (LabDiet), a standard natural ingredient diet containing 1.9 mg/kg folic acid, and 9.0 mg/kg riboflavin. In mice, based on numbering from ENSMUST00000069604.15, the 806C>T causes a codon change GCC>GCT, leading to an A262V mutation in the methylenetetrahydrofolate reductase gene that is equivalent to the 677C>T polymorphism and corresponding A222V mutation in humans. Given its common usage, we refer to this mouse model as *Mthfr^677C>T^*. Initially, the *Mthfr^677C>T^* was engineered by CRISPR onto LOAD1 (B6.*APOE4.Trem2*R47H*)^41^ (JAX #30922). To create the *Mthfr^677C>T^* strain (JAX #31464), *APOE4* and *Trem2*R47H* were bred out by successive backcrosses to B6. To produce experimental animals, mice heterozygous for *Mthfr^677C>T^* (*Mthfr^C/T^*) mice were intercrossed to produce litter-matched male and female *Mthfr^C/C^, Mthfr^C/T^* and *Mthfr^T/T^* mice (herein referred to as CC, CT and TT). Mice were aged to either 3-4 months, 6 months, 12 months, 18 months and 24 months.

### Mouse Perfusion, Tissue Preparation and Sectioning

Mice were anesthetized with a lethal dose of ketamine/xylazine based on animal mass and transcardially perfused with 1X PBS (Phosphate Buffered Saline). Liver blanching was used to determine a successful perfusion. Livers were removed for enzyme activity assessment and snap frozen in liquid nitrogen. Brains were dissected and hemisected at the midsagittal plane.

For enzyme activity, and assessment and metabolite assessment, brain hemispheres were snap frozen in liquid nitrogen. For immunohistochemistry assessment, brain hemispheres were fixed in 4% Paraformaldehyde (PFA) overnight at 4°C followed by 24 hours in 15% sucrose/Phosphate Buffered Solution (PBS) and 24 hours in 30% sucrose/PBS solution. Tissue was then frozen in OCT, and cryosectioned sagittally at 20 μm.

### Enzyme Activity and Western blotting

#### MTHFR enzyme activity

Liver and brain were collected from male mice (n=6/genotype except for TT; n=4) and from female mice in a separate cohort (n=6/genotype). MTHFR activity was determined using ^14^C-methylTHF-menadione oxidoreductase assay^42^ ( 37°C, reverse direction) with the following modifications: ^14^C-methylTHF was lowered to 200 μM, and total volume increased to 279 μL. Assays were performed in duplicate with 1 blank per sample, using 200 μg protein (from 10 μg/μL aliquots), by an individual blinded to genotype.

#### Protein extraction

Crude protein extract was prepared from 80 – 100 mg of frozen liver, (*n*=6/sex/genotype, except TT males; *n*=4) and whole half brain (*n*=6/genotype) in extraction buffer (50 mM potassium phosphate pH 7.2, 50 mM sucrose, 0.3 mM EDTA containing protease and phosphatase inhibitors (Pierce Protease Inhibitor Tablet and Halt Phosphatase Inhibitor Cocktail, ThermoScientific catalog numbers A32955 and 78420)) using a bead mill with steel beads (2 x 2 min at 20 Hz; TissueLyser, Qiagen). Extracts were cleared by centrifugation (17,000 RCF) at 4°C for 30 min. Protein content was determined by Bradford assay using BSA (Bovine Serum Albumin) as a standard. A portion of the extract was diluted to 10 μg/μL in extraction buffer and aliquoted. Diluted aliquots and concentrated extracts were stored at −80°C.

#### MTHFR stability assays

MTHFR stability was assessed by pre-incubating 800 μg of protein extract (from 10 μg/μL aliquots) at either 37 or 46°C in the MTHFR assay mix lacking cofactors (i.e. FAD and methylTHF). After 10 mins of incubation, ^14^C-methylTHF and FAD were added to the reaction mix, and the assay proceeded as described^42^. Assays were performed by an individual blinded to genotype, in duplicate for each temperature, with one blank at 37°C, as temperature was pre-determined not to affect ^14^C counts in the blank. (Males only, *n* = 3/genotype)

#### Western blotting

Western blots were performed with 25 μg of the protein extracted for MTHFR assays, using primary antibodies specific for MTHFR^9^, and β-actin (A2066; Sigma-Aldrich), as in^43^. Blots were imaged using the Amersham Imager 600 and quantitated with the Amersham Imager 600 analysis software v1.0.0 (GE Healthcare Life Sciences). (Males *n*=6/genotype except for TT *n*=4; females *n*=6/genotype)

### Homocysteine

Blood was collected by cardiac puncture from non-fasted, anesthetized animals (see Perfusion method) at harvest prior to perfusion (*n*=8-12/sex/genotype/age) using a 25-gauge EDTA-coated needle, attached to a 1mL syringe. 300-500mL of blood was slowly injected into a 1.5mL EDTA coated MAP-K2 blood microtainer (363706, BD, San Jose, CA) at room temperature. Blood tubes were spun at 23°C and 4388 RCF for 15 minutes. Blood plasma was removed and aliquoted into 1.5mL tubes. Tubes were stored at −20°C before analysis. Thawed blood plasma was analyzed by Beckman Coulter AU680 chemistry analyzer (Beckman Coulter, Brea, CA) and Siemens Advia 120 (Germany) for levels of homocysteine. Human homocysteine values were obtained from a defined cohort Alzheimer’s Disease Neuroimaging Initiative (ADNI)

### Folate and methionine cycle plasma and brain tissue metabolites

Brain tissue folates were determined by liquid chromatography coupled with electrospray positive ionization tandem mass spectrometry (LC-ESI-MS/MS) as previously described^44^. Briefly, brain tissue (25 mg) was sonicated in 4 volumes of extraction buffer (HEPES buffer pH 8) to which 40 μL of charcoal treated rat plasma was add, as a source of pteroylpoly-γ-glutamate hydrolase for the deconjugation of the polyglutamate forms to monoglutamate forms of folates. Following centrifugation, filtering, and addition of ^13^C_5_-methyl-THF internal standard (IS) clear extracts were injected in the UHPLC-MS/MS system consisting of a Waters Acquity UPLC coupled to a Xevo-TQS triple quadrupole mass spectrometer equipped with an electrospray ionization probe (Waters Corporation, Milford, MA, USA). Quantification of folate forms was achieved using the peak area ratio (area of metabolite/area of IS).

Brain SAM, SAH, methionine, cystathionine, choline and betaine were determined by LC-ESI-MS/MS following sonication of tissue with 4 volumes of perchloric acid (0.4M) as previously described^45, 46^. Clear perchloric acid extracts containing labeled isotopes were injected into the LC-MS/MS system (Qtrap 5500, Sciex, Framingham, MA) and metabolites quantitated using Analyst 6.0 (Sciex, Framingham, MA). (n=6/sex/genotype except male TT; *n*=4).

### PET/CT

To assess neurovascular perfusion, mice were non-invasively imaged via PET/CT (*n*=5-6 mice/sex/genotype/age). PET/CT is used to give both anatomic (CT) and functional (PET) information, and were performed sequentially during each session, thus permitting the images to be co-registered for regional specificity analysis. To measure regional blood flow, 5.3±0.09 MBq (in 100 μL) copper-pyruvaldehyde-bis(N4-methylthiosemicarbazone) (^64^Cu-PTSM)^47^, which has a very high first pass (>75%) extraction^48^, and glutathione reductase redox trapping of copper^48^, was administered via tail vein in awake subjects and given a 2 min uptake period in their warmed home case prior to imaging. Post uptake, mice were induced with 5% isoflurane (95% medical oxygen) and maintained during acquisition with 1-2% isoflurane at 37°C. To provide both anatomical structure and function, PET/CT imaging was performed with a Molecubes β-X-CUBE system (Molecubes NV, Gent Belgium). For PET determination of tissue perfusion, calibrated listmode PET images were acquired on the β-CUBE, and reconstructed into a single-static image using ordered subset expectation maximization (OSEM) with 30 iterations and 3 subsets^49^. To provide anatomical reference, and attenuation maps necessary to obtain fully corrected quantitative PET images, helical CT images were acquired with tube voltage of 50 kV, 100 mA, 100 μm slice thickness, 75 ms exposure, and 100 μm resolution. In all cases, images were corrected for radionuclide decay, tissue attenuation, detector dead-time loss, and photon scatter according to the manufacturer’s methods^49^. Post-acquisition, all PET and CT images were co-registered using a mutual information-based normalized entropy algorithm^50^ with 12 degrees of freedom, and mapped to stereotactic mouse brain coordinates^51^. Finally, to quantify regional changes, voxels of interest (VOI) for 27 brain (54 bilateral) regions were extracted and analyzed for SUVR (relative to cerebellum) according to published methods^52^.

### Autoradiography

To confirm the *in vivo* PET images, and to quantify regional tracer uptake, brains were extracted post rapid decapitation, gross sectioned along the midline, slowly frozen on dry ice, embedded in cryomolds with Optimal Cutting Temperature (OCT) compound (Tissue-Tek), and 20μm frozen sections were obtained at prescribed bregma targets (n = 6 bregma/mouse, 6 replicates/bregma) based on stereotactic mouse brain coordinates^51^. Sections were mounted on glass slides, air dried, and exposed on BAS Storage Phosphor Screens (SR 2040 E, Cytiva Inc.) for up to 12 h. Post-exposure, screens were imaged via Typhoon FL 7000IP (GE Medical Systems) phosphor-imager at 25μm resolution along with custom 64Cu standards described previously^53^.

### Immunostaining

20 μm cryosections were rinsed once in 1X PBT (PBS + 1% Triton 100X) and incubated in primary antibodies diluted with 1X PBT + 10% normal donkey serum overnight at 4°C. After incubation with primary antibodies, sections were rinsed three times with 1X PBT for 15 min and incubated overnight at 4°C in the corresponding secondary antibodies. Tissue was then washed three times with 1X PBT for 15 min, incubated with DAPI for 5 minutes, washed with 1X PBS and mounted in Poly aquamount (Polysciences). The following primary antibodies were used: goat anti-Collagen IV (ColIV) (1:40, EMD Millipore), chicken anti-GFAP (1:200, Origene). Secondary antibodies: donkey anti-goat 488 (1:500, Invitrogen) donkey anti-chicken 594 (1:500, Jackson ImmunoResearch Laboratories).

### Confocal imaging and Imaris Analysis

Z-stacks of immunostained brain sections were taken on a confocal microscope (Leica TCS Sp8 AOBS with 9 fixed laser lines) at 20x magnification using the Leica Application Suite X (LAS X) software (Leica Microsystems GmbH, Mannheim, Germany). Three images were taken for each brain from each mouse from 1) frontal cortex, 2) somatosensory cortex and 3) visual cortex (*n*=6/sex/genotype/age for a total of 216 images). Imaris software (x64 9.5.1) was used for quantification of ColIV^+^ vessel area as well as area covered by GFAP^+^ astrocytes in the cortex, using a workflow developed in-house, available upon request. Briefly, confocal images were converted into Imaris compatible files and vessel or astrocyte boundaries were confirmed. For each image, individual vessel or astrocyte volumes were determined in μm^3^ and collated in Excel. The sum of the individual vessel or astrocyte volumes were determined per image, and this value was used for statistical analysis.

### Transmission Electron Microscopy (TEM)

Mice were perfusion fixed (using 2% PFA and 2% glutaraldehyde in 0.1 M Cacodylate buffer (Caco)). Brains were fixed overnight in the same solution at 4°C then stored in 1X PBS. 100 μm thick sagittal sections were cut on a vibrating microtome and post fixed with 2% osmium tetroxide in 0.1 M Caco buffer for 2 hours at room temperature. Sections were rinsed three times with Caco buffer and then dehydrated through of series of alcohol gradations. The sections were then put into a 1:1 solution of propylene oxide/Epon (EMED 812) overnight on an orbital rotator. Sections were then flat embedded with 100% Epon between 2 sheets of Aclar film and polymerized at 65°C for 48 h. For each section, a region of the frontal cortex was cut out and glued onto dummy blocks of Epon. 90 nm ultrathin sections were cut on a Diatome diamond knife, collected on 300 mesh copper grids, and stained with Uranyl Acetate and lead citrate. Grids were viewed on a JEOL JEM1230 transmission electron microscope and images collected with an AMT high-resolution digital camera. For each brain, 10-20 cross-sectional blood vessels as well as other defining cells or features (e.g., microglia) were imaged per (*n* = 3/sex/genotype.) In a blinded analysis, images were assessed for the following main features 1) presence/absence of endothelial cells, 2) open/closed lumen, 3) presence/absence of astrocyte endfeet, 4) presence/absence of apoptotic cells, 5) presence/absence of microglia and if present, their level of interaction with vessels.

### Statistical Analysis

Data are shown as the mean ± standard deviation. Multiple group comparisons were performed by one-way or two-way multifactorial analysis of variance (ANOVA) depending on the number of variables followed by Tukey post hoc test. When assessing differences between two groups, a paired two-tailed Student’s test was used. Data from PET/CT were assessed in two ways: principal component analysis as a data reduction method, where regions explaining 80% of the variance were selected for statistical analysis, and by multivariate analysis of variance (MANOVA). Correlation and linear regression analyses were performed when assessing MTHFR activity vs protein expression and when assessing BMI vs plasma homocysteine levels. Differences were considered to be significant at *p* < 0.05. Statistical analyses were performed utilizing Prism v7.0 h software (GraphPad Software Inc., San Diego, CA, USA) or SPSS 28.0.1 software. For all analyses, * p<0.05, ** p<0.01, *** p<0.001, **** p<0.0001.

## Results

### *Mthfr^677C>T^* mice display reduced enzyme activity in liver, with relevant metabolic changes in plasma

Since liver is a major tissue for folate metabolism, MTHFR enzyme activity was assessed in crude liver extracts of 3-4 month old male and female *Mthfr^677C>T^* mice. In both sexes, enzymatic activity was significantly reduced in mice carrying one (CT, heterozygous) or two (TT, homozygous) copies of the risk allele compared to wild type mice (CC). Although specific activity did not differ significantly between the CC and CT groups, activity in the TT mice was significantly lower than both the CC and CT groups (**Figure 1a**). These data are consistent with human findings conducted in human leukocytes^9^. Presence of the variant increased the thermolability of human MTHFR, so thermostability was assessed in crude liver extracts of male *Mthfr^677C>T^* mice. Percent retained activity in CC, CT and TT liver extracts decreased significantly in a genotype-specific manner. Specifically, in Tukey post-hoc comparisons, CT was less stable than CC, and TT was significantly less stable than both CC and CT (**Figure 1b**).

**Figure 1.**
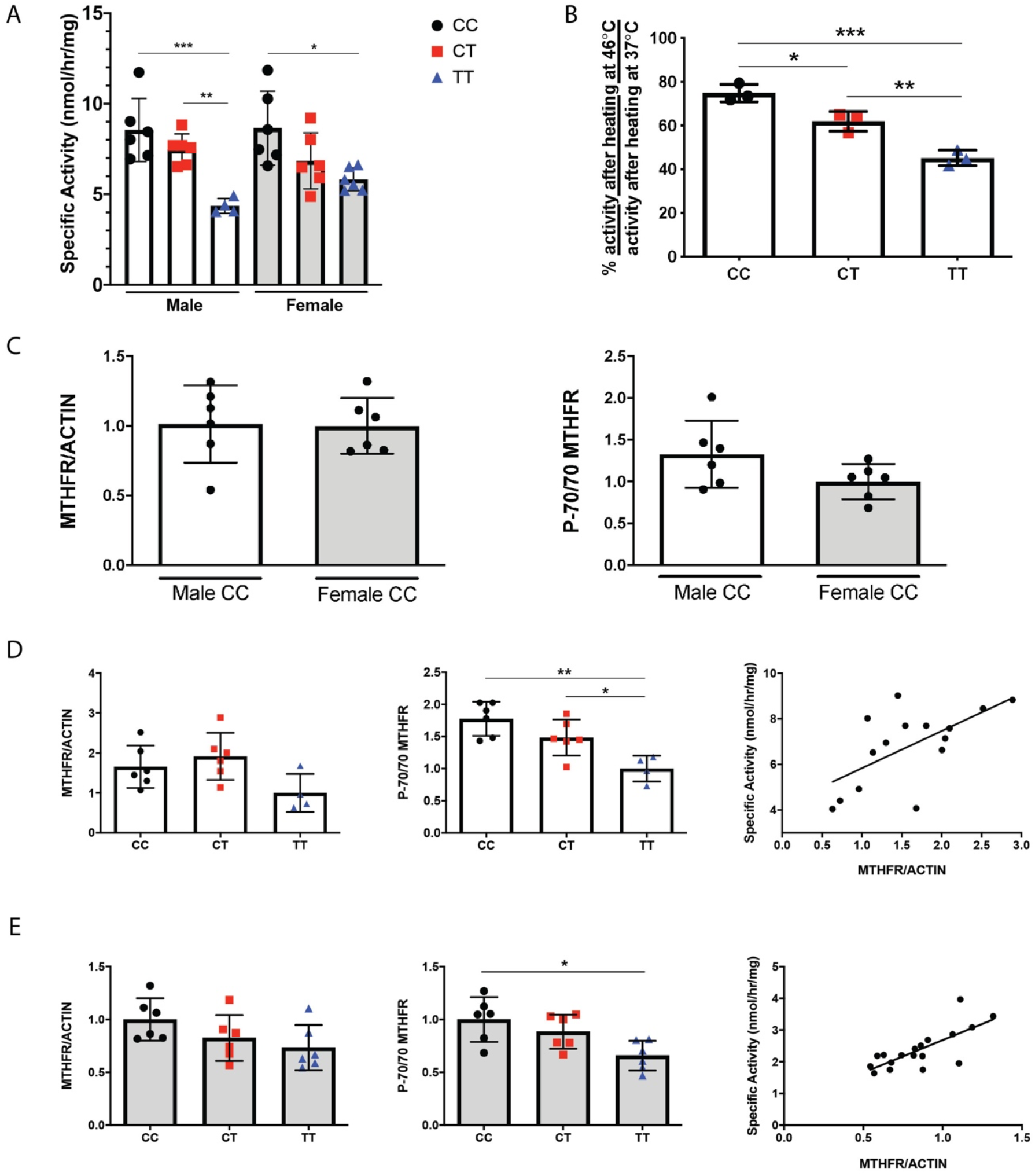
(**A**) MTHFR enzymatic activity in the liver. Activity is significantly reduced in both males and females carrying one (CT) or two (TT) copies of the variant. (n=6/genotype except for TT males: n=4; P=0.0002 One-way ANOVA, Tukey post-hoc). (**B**) Liver enzymatic activity after the sample is heated is shown as a ratio of percent activity at 46 degrees Celsius/percent activity at 37 degrees Celsius. Activity is significantly reduced in male CT and TT mice. (n=3/genotype; P=0.0003, One-way ANOVA, Tukey post-hoc). (**C**) Western blots were performed on control (CC) male and female mice and determined that MTHFR protein expression does not differ by sex (left) using actin as a standard (n=6/sex; unpaired t-test). The ratio of the phosphorylated MTHFR isoform p70 to the non-phosphorylated isoform is also not significantly different between male and female controls (right; n=6/genotype; unpaired t-test). (**D**) Within males, total MTHFR does not significantly differ based on genotype (left; n=6/genotype except for TT males: n=4; One-way ANOVA, Tukey post-hoc). However, the ratio of phosphorylated to non-phosphorylated isoforms is reduced in CT and TT males (center; n=6/genotype except for TT males: n=4; P=0.0017, One-way ANOVA, Tukey post-hoc). MTHFR enzymatic activity correlates with total MTHFR protein (right; n=6/genotype except for TT males: n=4; correlation with activity, P=0.0127 Pearson r=0.6250). (**E**) In females, total MTHFR protein also does not significantly differ based on genotype (left; n=6/genotype; One-way ANOVA, Tukey post-hoc). The ratio of phosphorylated to non-phosphorylated isoforms is reduced in TT females (center; n=6/genotype; P=0.0119, One-way ANOVA, Tukey post-hoc). MTHFR enzymatic activity correlated with total MTHFR protein (right; n=6/genotype; correlation with activity, P= 0.0004 Pearson r=0.7471).

MTHFR protein levels were assessed in liver extracts by Western blot. There was no significant effect of sex on MTHFR levels, (**Figure 1c**) as was previously reported^54^. Similarly, there was no significant effect of sex on the ratio of the phosphorylated to non-phosphorylated isoforms of MTHFR (**Figure 1c**). MTHFR levels did not differ significantly due to genotype (**Figure 1d-e**). The phosphorylated (P-70) isoform of MTHFR is reported to be less active than the nonphosphorylated (70) isoform^55, 56^. The ratio of the P-70 to 70 isoforms of MTHFR differed significantly between genotypes and was significantly lower in the TT livers of both males and females when compared to both CC and CT due to the increase in the non-phosphorylated isoform. It is possible that this shift towards a more active form may be a compensatory response to the reduced expression/activity of MTHFR (**Figure 1d and 1e**). There was good correlation between MTHFR activity and expression (**Figure 1d, 1e**), which suggests that the reduced MTHFR activity observed in the mouse model and in people with the TT genotype^9, 42^, may be due to reduced amounts of MTHFR protein in liver and possibly in other tissues.

With less available MTHFR enzyme activity to convert homocysteine into methionine, more homocysteine remains in the blood and has been shown to promote vascular inflammation^57–59^. Therefore, plasma homocysteine levels were assessed in *Mthfr^677C>T^* mice at four ages: 6, 12, 18 and 24 months old (**Figure 2a**). Notably, 6 month old TT mice showed the most distinct increase in homocysteine levels compared to littermate controls (**Figure 2a**). Older groups demonstrated sex differences (female mice have overall higher homocysteine levels) but not significant differences between genotypes. However, within older female mice carrying the variant, homocysteine levels still trended at an elevated level. Human data from the Alzheimer’s Disease Neuroimaging Initiative (ADNI) showed that overall, subjects within the cohort presented with higher plasma homocysteine based on *MTHFR^677C>T^* genotype (**Figure 2b**). Unlike mice, male humans showed higher overall plasma homocysteine levels when the data are stratified by sex (**Figure 2c**). This discrepancy may be diet-related as men in the ADNI study had higher overall Body Mass Index values, (**Figure 2d**) while mice in these studies were fed a standard low-fat chow for the duration of their lives and did not become obese as they aged.

**Figure 2.**
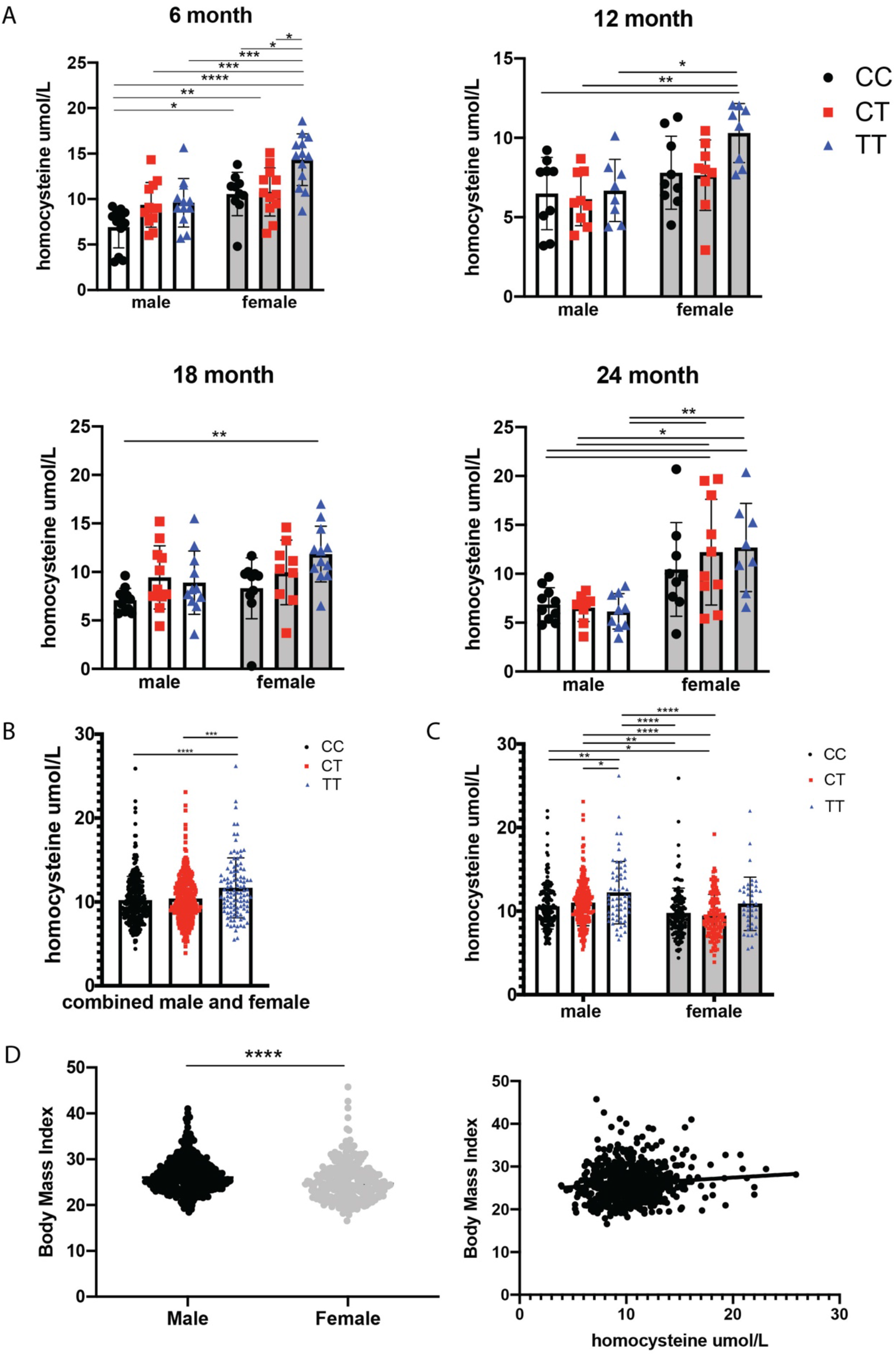
(**A**) Plasma homocysteine levels in *Mthfr^C677T^* mice at 6, 12, 18 and 24 months of age. At 6 months, homocysteine levels are significantly elevated in a genotype-specific manner. At 12, 18 and 24 months, female mice have significantly elevated homocysteine compared to males within the same age group. (*n*=8-12/sex/genotype/age; Two-way ANOVA, Tukey post-hoc for each age group. 6 month P=<0.0001 for sex, P=0.0002 for homocysteine, 95% CI {-4.494 to −2.058} 12 month P=0.0005 for sex, 95% CI {-3.302 to −0.9893} 18 month P=0.0379 for sex P=0.0114 for homocysteine, 95% CI {-3.031 to 0.09010} 24 month P=<0.0001 for sex, 95% CI {-7.242 to −3.255}). (**B**) Data from a human cohort (ADNI) demonstrates a similar trend in genotype-driven homocysteine elevation (left; n=442 males, 296 females: P=<0.0001, One-way ANOVA, Tukey post-hoc). (**C**) When males and females are separated, males drive the trend (right; P=<0.0001 for both sex and genotype, Two-way ANOVA, 95% CI {0.7415 to 1.706}, Tukey post-hoc). (**D**) Overall, males within the cohort have a higher body mass index (BMI) than females (left; n=369 males, 259 females: P=<0.0001, unpaired t-test) and there is a modest but significant correlation between homocysteine and BMI within the cohort (right; correlation with activity, P=0.0095 Pearson r=0.1034).

Because plasma homocysteine levels showed genotype-dependence, plasma from 3-4 month old mice was assessed for metabolites within the folate and methionine pathways. DHF (dihydrofolic acid) is a folic acid derivative that is converted to THF (tetrahydrofolic acid) by dihydrofolate reductase. One-carbon substituted THFs are cofactors in a number of reactions, including the synthesis of amino acids and nucleic acids and can serve as carriers for methyl groups. MTHFR generates 5-methylTHF (5-MTHF), which participates in the remethylation of methionine and is further reduced to THF. Betaine, a choline metabolite, can also remethylate homocysteine to methionine. S-adenosyl methionine (SAM) and S-adenosyl homocysteine (SAH) are involved in methyl group transfers. Cystathionine is an intermediate in the synthesis of cysteine from homocysteine. Interestingly, only cystathionine showed a significant difference based on genotype, with female TT mice showing elevated levels compared to CC (**Supplementary Figure 1**). This result is consistent with higher levels of homocysteine in female TT mice. Collectively these assays provide early characterization of the new model and demonstrate the similarity between the *Mthfr^677C>T^* mouse model and human carriers of *MTHFR^677C>T^*.

### *Mthfr^677C>T^* reduces enzyme activity and alters metabolic profiles in brain

Historically, the effects of null mutations in *Mthfr* have been studied systemically, but we were interested in examining the role of the 677C>T variant in the brain, hypothesizing that the immune-privileged central nervous system could be more susceptible to variant-induced damage. Published brain RNA-seq data show *Mthfr* is expressed at low levels in numerous cell types in mouse brain, with the highest expression being in microglia/macrophages and endothelial cells in adult mice (**Figure 3a**)^60^. Similarly, available single-cell RNA-seq data show *Mthfr* is expressed in the majority of mouse brain cell types assessed but most strongly in arterial smooth muscle cells (aSMC) and a subset of endothelial cells (**Figure 3b**)^61, 62^.

**Figure 3.**
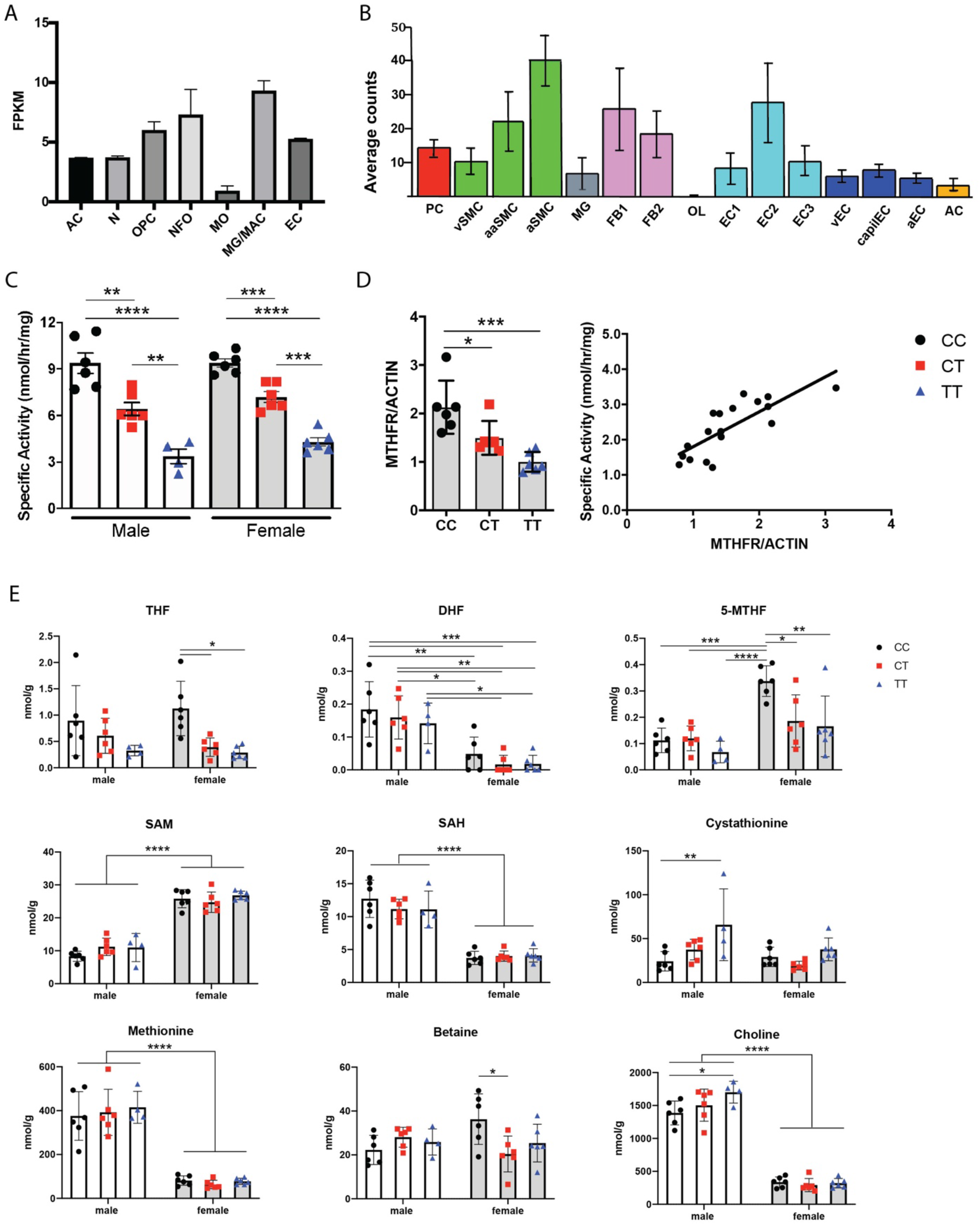
(**A**) Numerous cell types in the brain express *Mthfr*. Data obtained from the Brain RNA-seq database. (**B**) Single cell sequencing showing expression of *Mthfr* in cerebrovascular and perivascular cells. Data obtained from the Single-Cell RNA-seq Gene Expression Data. AC= astrocytes, N= neurons, OPC= oligodendrocyte precursor cells, NFO= newly formed oligodendrocytes, MO= myelinating oligodendrocytes, MG= microglia, MAC= macrophages, EC= endothelial cells, PC= pericytes, vSMC= venous smooth muscle cells, aaSMC= arteriolar smooth muscle cells, aSMC= arterial smooth muscle cells, FB1= fibroblast-like type 1, FB2= fibroblastlike type 2, OL= oligodendrocytes, EC1= endothelial cell type 1, EC2= endothelial cell type 2, EC3= endothelial cell type 3, vEC= venous endothelial cells, capilEC= capillary endothelial cells, aEC= arterial endothelial cells. (**C**) MTHFR enzymatic activity measured in brain tissue. Both males and females show significantly reduced activity based on genotype (n=4-6/sex/genotype; P=<0.0001, One-way ANOVA, Tukey post-hoc). (**D**) In brains of female mice, total MTHFR is significantly reduced in both the CT and TT groups (left; n=6/genotype; P=0.0002, One-way ANOVA). Enzymatic activity is correlated with MTHFR protein expression (right; correlation with activity, P=0.0001 Pearson r=0.7896). (**E**) Brain-derived metabolite levels: THF= tetrahydrofolic acid, DHF= dihydrofolic acid, 5-MTHF= 5-methyltetrahydrofolate, SAM= S-Adenosyl methionine, SAH= S-Adenosylhomocysteine. DHF (P= <0.0001; 95% CI {0.09381 to 0.1738}), 5-MTHF (P= <0.0001; 95% CI {-0.1831 to −0.07584}), SAM (P= <0.0001; 95% CI {-17.52 to −13.76}), SAH (P= <0.0001; 95% CI {6.445 to 8.966}) methionine (P= <0.0001; 95% CI {270.3 to 369.9}) and choline (P= <0.0001 95% CI {1107 to 1324}) demonstrate significant sex-differences, while THF (P= 0.0007; 95% CI {-0.2728 to 0.2883}), 5-MTHF (P= 0.0075; 95% CI {-0.1831 to −0.07584}), cystathionine (P= 0.0031 95%CI {1.798 to 25.69}) and betaine (P= 0.0086 95% CI {-7.633 to 3.749}) show genotype-specific differences (n=4-6/sex/genotype; Two-way ANOVA, Tukey post-hoc).

MTHFR activity in brains from 3-4 month old mice demonstrated a similar genotype-based reduction in activity as seen in liver, in both males and females (**Figure 3c**). Brain extracts from 3-4 month old female *Mthfr^677C>T^* mice were assessed by Western blot using actin as a loading control. Unlike data from the liver, MTHFR protein expression was reduced in a genotype-specific manner in the TT brain tissue, suggesting that the 677C>T variant may produce enhanced effects in the central nervous system compared to the liver. Again, there is good correlation between MTHFR activity and expression (**Figure 3d**). Brain tissue from 3-4 month old mice was also assessed for metabolites within the folate and methionine pathways. Unlike the parallel metabolite data from plasma (**Supplementary Figure 1**), THF and 5-MTHF were significantly reduced in brain tissue of female CT and TT mice, and betaine was significantly reduced in CT females compared to CC controls. Cystathionine was significantly elevated in male TT mice, (**Figure 3e**) which mimics the female plasma cystathionine results (**Supplementary Figure 1**). Interestingly, results for DHF, SAM, SAH, methionine and choline showed significant sex-differences, with females presenting with lower levels for all of these metabolites except for SAM (**Figure 3e**).

Together, these data show that *Mthfr^677C>T^* decreases MTHFR protein expression and enzyme activity in the brain suggesting it may play distinct roles in the brain and periphery.

### *Mthfr^677C>T^* promotes reduced cerebral blood perfusion

Because we observed metabolic changes in the Mthfr^677C>T^ brain, plasma and liver, we next determined whether these effects were sufficient to induce cerebrovascular deficits. In human studies the 677C>T variant is associated with increased risk for vascular inflammation and alterations to blood flow. Therefore, to quantify the regional changes in brain perfusion, we performed *in vivo* PET/CT imaging using ^64^Cu-PTSM, and autoradiography (autorad) was used to confirm the findings (**Figure 4a**). Two statistical approaches – principal component analysis (PCA) and MANOVA – were used to identify significant changes in blood flow, which collectively showed differences based on genotype. PCA compared sex, genotype and age and determined thirteen brain regions that explained 80 percent of the variance in blood flow in all mice and all conditions (**Figure 4b**). Results demonstrating specific regional statistics by MANOVA are listed in **Supplementary Table 1**. Interestingly, 6 month old mice showed reduced perfusion overall compared to 18 month old mice. While some regions showed a reduction in blood perfusion within both CT and TT mice, a number of regions showed a prominent reduction in the CT group only (**Figure 4b**). Several regions that showed reduced blood perfusion in a genotype-dependent manner are involved in memory, learning, sensory and motor function.

**Figure 4.**
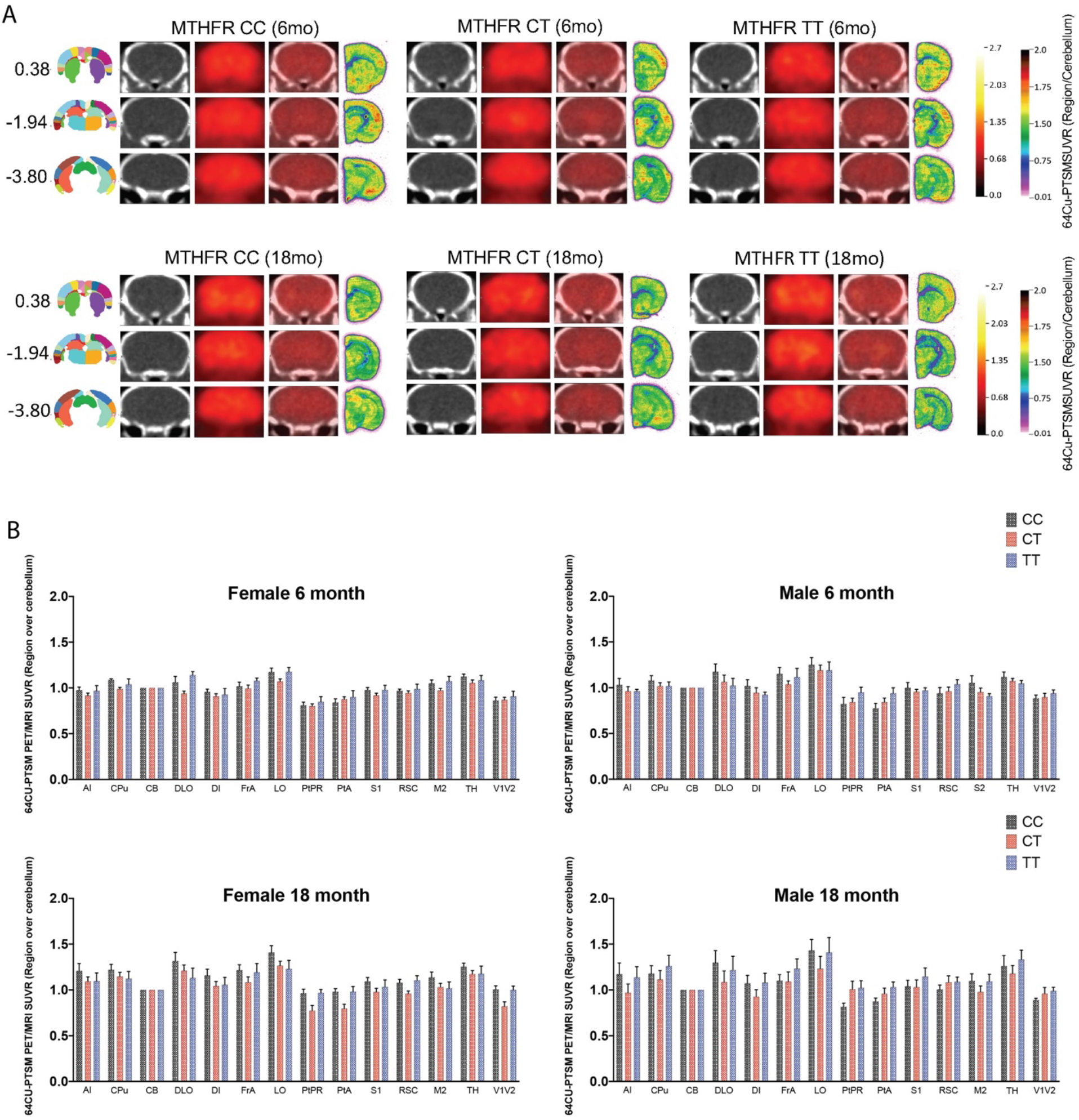
(**A**) Representative images are shown from mice at 6 and 18 months of age. From right to left, the anatomical location of the image as determined by the Allen Mouse Brain Atlas, CT images showing clear locations of identifiable brain structures, PET images showing changes in blood perfusion via tracer concentration, the overlay of the PET and CT, and finally autorad demonstrating radioactive decay on the far right (n=6/sex/genotype/age). (**B**) 13 brain regions were determined by principal component analysis to explain 80% of the variance in blood flow within all mice. Data is presented as brain regions normalized to the cerebellum.

### Immunohistochemistry and electron microscopy reveal cerebrovascular deficits in *Mthfr^677C>T^* mice

Given the expression of *Mthfr* in multiple vascular related cells including endothelial and smooth muscle cells, and the reduction in blood flow by PET, two key components of the BBB were assessed by immunohistochemistry. Examination of endothelial-associated basement membrane protein Collagen IV (ColIV) and the astrocyte marker GFAP in the frontal cortex, somatosensory cortex and visual cortex revealed genotype-dependent phenotypes (**Figure 5**). Analysis of ColIV volume per image demonstrated that vascular density was not significantly altered in the visual or somatosensory regions due to sex or age, but a trend of reduced density was observed in both CT and TT mice. However, ColIV+ vascular density was significantly reduced in the frontal cortex of 6-month old CT and TT mice but did not reach significance at 18 months (**Figure 5b**). Analysis of GFAP+ astrocyte volume per image showed a significant increase in astrocyte density in the visual and frontal cortical regions of TT males, with a genotype-specific trend in 6 month old females (**Figure 5c**). All 18 month old groups demonstrated higher numbers of GFAP-expressing astrocytes at baseline, but also showed a genotype-specific trend, suggesting increased astrocyte reactivity in regions with lower vascular density.

**Figure 5.**
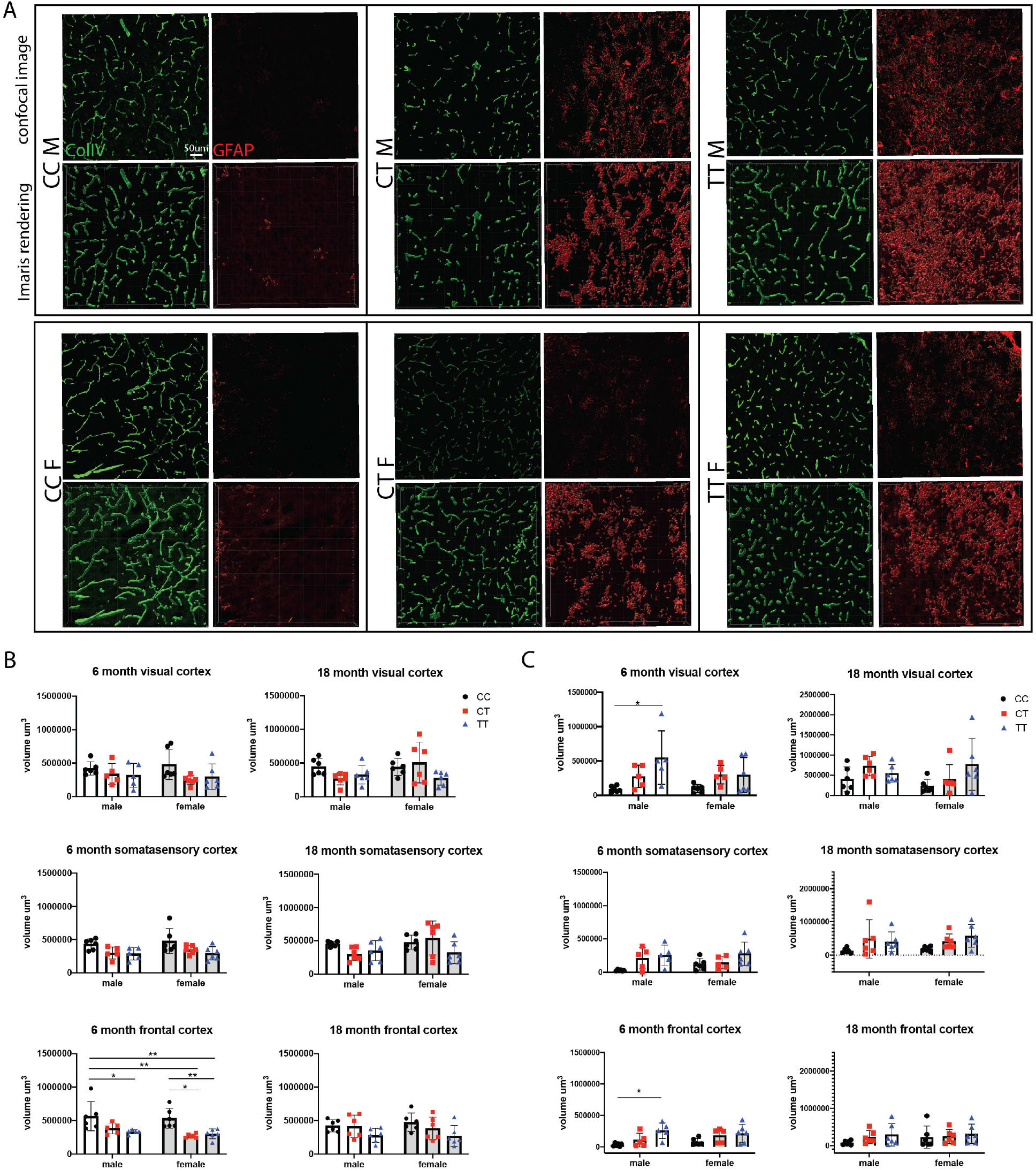
(**A**) From right to left, representative confocal images of immunolabeled Collagen IV-expressing vascular basement membrane, GFAP-expressing astrocytes, and their respective Imaris renderings in 6 month male and female mice from each genotype. (**B**) Vascular density was assessed in 3 regions of the cortex at 6 and 18 months of age. The volume of ColIV per image was calculated using Imaris software and the sum of the volume/image was used for analysis. 6 month old males and females show significantly reduced vascular volume in the frontal cortex of TT or both CT and TT mice, respectively (n=6/sex/genotype/age; P=<0.0001 for genotype, 95% CI {-36337 to 89790} Two-way ANOVA, Tukey post-hoc). (**C**) Astrocyte expression of GFAP was assessed in the same set of images using the same protocol for Imaris analysis. In 6 month old males, there was a significant increase in GFAP-expressing astrocytes in both the visual cortex and the frontal cortex (n=5-6/sex/genotype/age; visual cortex P=0.0026 95% CI {-73800-220053}, frontal cortex P=0.0012 95% CI {-98179 to 48259}, Two-way ANOVA, Tukey post-hoc).

Vascular morphology was further assessed in cross sections of cortical cerebrovasculature from *Mthfr^677C>T^* mice at 6 months of age using electron microscopy (**Figure 6**). Overall, CC mice of both sexes demonstrated healthy vasculature, including presence of endothelial cells surrounding an open lumen. CT and TT mice, however, often showed vessels with a closed or missing lumen, possibly because of either cell loss or a thickened basement membrane or tunica intima. Additionally, we identified numerous instances of degenerating cells (likely either endothelial cells or pericytes identified by morphology) with an apoptotic phenotype. Furthermore, balloon-like structures that stained only faintly with osmium tetroxide often encircled vessels and are characteristic of swollen astrocytic endfeet^63^. These features were frequently associated with an increased number of activated microglia/monocytes, characterized by high electron density, large rough endoplasmic reticulum, and visibly engulfed cell components, suggesting that these microglia/monocytes were phagocytic when near vasculature. While some samples from mice with CT or TT genotypes contained only one notable abnormality, most contained multiple abnormalities.

**Figure 6.**
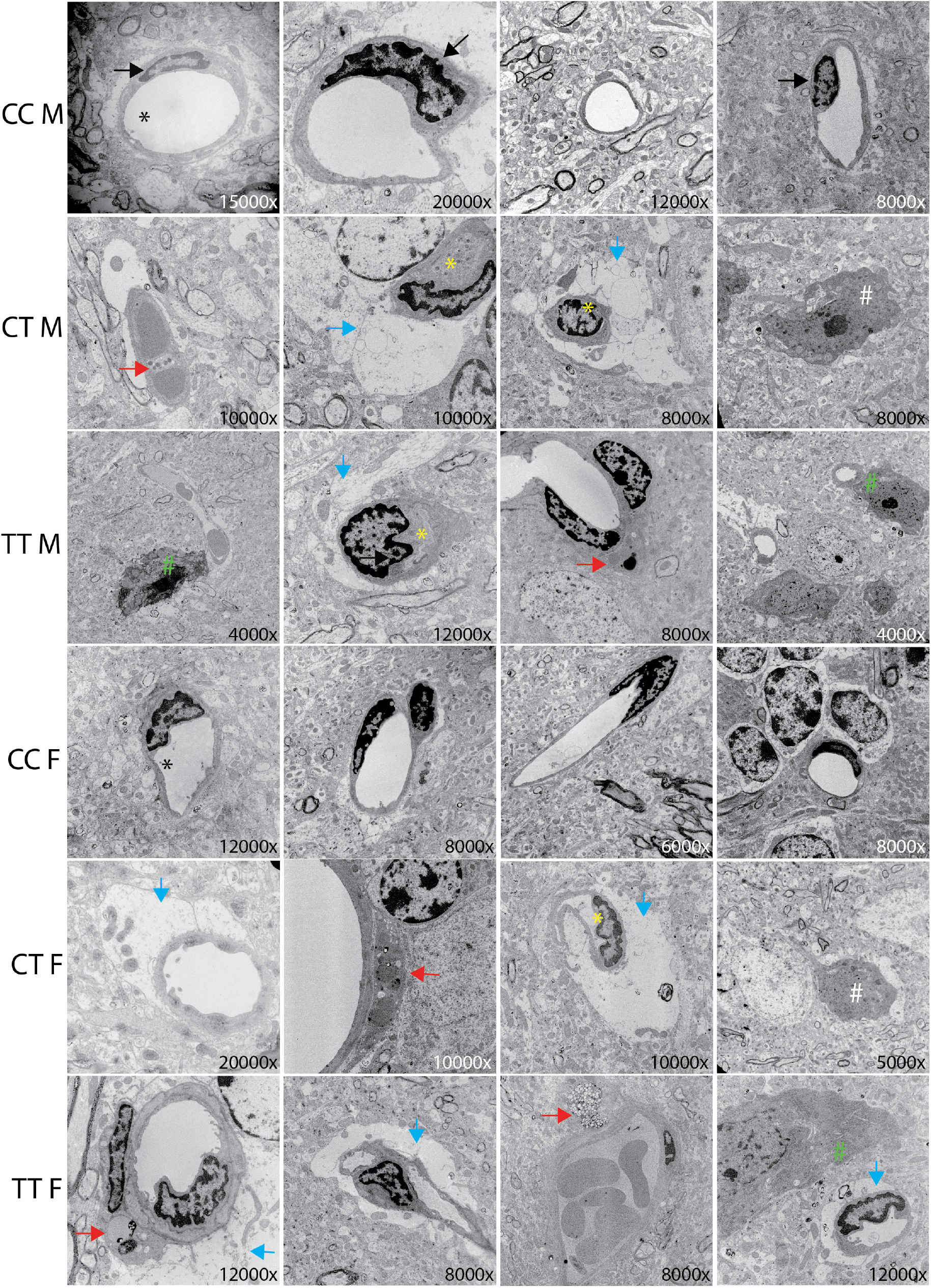
Cerebrovascular ultrastructure in cortex is visualized using electron microscopy. Images are representative of typical vascular features from each genotype. Magnification is noted in the bottom right corner of each image. Relevant features are noted in the figure by arrows, asterisks, and hashtags including: Black arrow= endothelial cell, Black asterisk= open lumen, yellow asterisk closed lumen, red arrow= cells going through apoptosis, blue arrow= swollen astrocyte endfeet, white hashtag= significant presence of microglia, green hashtag= microglia interacting with a vessel (n=3/sex/genotype).

## Discussion

With an increased focus on vascular contributions to dementias, there is a clear need to increase the number of human-relevant genetic-based mouse models that recapitulate endophenotypes of VCID. In this study we demonstrated that *Mthfr^677C>T^* mice phenocopy outcomes observed in humans carrying *MTHFR^677C>T^* and identify novel cerebrovascular phenotypes. Depending on ethnicity and geographic location, up to 40 percent of people can be heterozygous and up to 20 percent can be homozygous for the T risk allele^14^. Therefore, it is likely that the 677C>T variant alone does not promote pathology that directly results in cognitive decline. Baseline cerebrovascular dysfunction may decrease resilience and increase susceptibility when additional genetic and/or environmental risk is introduced. In order to better understand disease initiation and progression, it is critical to consider how multiple systems work in tandem to generate pathology. To this end, MODEL-AD is focused on combining known variants such as *APOE^E4^, TREM2^R47H^* and *Mthfr^677C>T^* and introducing alternative diets commonly consumed in the western world.

Recapitulating human data, we observed MTHFR enzymatic deficiency in the liver as well as an increase in plasma homocysteine. Total MTHFR protein in the liver was not significantly different based on genotype, but the ratio of the inactive phosphorylated MTHFR isoform (P-70) to the activated non-phosphorylated isoform (70) was genotype-dependent. We speculate that the activated form may be increased in TT mice to compensate for the reduction in enzyme activity. Interestingly, homocysteine levels were significantly elevated in the 6 month old female CT and TT mice but this was not sustained with age. Similar effects have been reported in human studies^64^. It is possible that *Mthfr^677C>T^* exerts its greatest effect during development or early life and compensatory mechanisms appear with age. Female mice consistently had higher homocysteine levels than males, regardless of genotype or age. However, data from the ADNI human cohort suggests that in humans, men have slightly higher levels of homocysteine overall as well as significant genotype specific homocysteine elevation. When BMI was stratified by sex, we see that men within the cohort have higher overall BMIs than women. *Mthfr^677C>T^* mice are fed a balanced diet with low-fat composition, while diet is not accounted for in the ADNI cohort. It is plausible that diet and corresponding weight changes may have an effect on plasma homocysteine, but future studies will be required to make this assertion.

Historically, *MTHFR* polymorphisms (677C>T and 1298A>C) have been studied in the context of systemic and reproductive dysfunction^65, 66^. Mild to moderate hyperhomocysteinemia has been previously implicated as a risk factor for hypertension^67^ and cardiovascular disease^68^. Presence of these variants, with and without mildly elevated levels of homocysteine, has been studied in relation to several multifactorial disorders including recurrent pregnancy loss^69^, neural tube defects and congenital anomalies^70^, cancer^71^, glaucoma, and neurodevelopmental disorders, notably autism^72^. However, because the 677C>T variant has been linked to AD or vascular dementia in multiple GWAS, we hypothesized that it is playing a role specifically in the brain.

Compared with controls, MTHFR activity in the brain is significantly reduced in *Mthfr^677C>T^* mice, with activity being even lower than activity in the liver. Additionally, we observed a significant decrease in total MTHFR protein as well as genotype-specific reduction of folate and methionine cycle metabolites in female CT and TT mice. These metabolites were not significantly altered in circulating blood plasma from the same animals, suggesting a brain-specific sensitivity to metabolic changes induced by the variant. MTHFR deficiency may have more adverse effects in brain because the choline-betaine dependent homocysteine remethylation pathway has reduced activity in brain than in liver, thereby potentially comprising methylation capacity. In addition, this variant is stabilized by folate; since liver has greater folate stores than brain, MTHFR protein from T variant carriers may be less susceptible to degradation in liver compared to brain. Unexpectedly, sex differences in levels of DHF, SAM, SAH, methionine and choline were dramatically different regardless of genotype. However, clinical studies have shown that sex-dependent metabolic changes occur in serum and cerebral spinal fluid of AD patients^73, 74^ so disparities in this study may represent an unexplored avenue for sex-dependent folate and methionine cycle metabolism in the brain.

*Mthfr^677C>T^* mice demonstrated brain region-specific reduction in perfusion including the Dorsolateral Orbital Cortex (DLO), Lateral Orbital Cortex (LO) and Dysgranular Insular Cortex (DI). The DLO is important for executive functions including working memory, cognitive flexibility, planning, inhibition, and abstract reasoning. The LO is part of the prefrontal cortex that has extensive connections with sensory areas as well as limbic system structures involved in emotion and memory. The DI is one of the more complex anatomical hubs and mediates a wide variety of brain functions including emotion regulation, learning and memory and social interactions. Therefore, perturbations of tissue perfusion in *Mthfr^677C>T^* mice is likely to increase susceptibility to a decline in ADRD-relevant cognitive functioning. The notion that a change in perfusion is altered by the introduction of a single variant emphasizes the role of *Mthfr^677C>T^* in regulating vascular function and underlines the concept of additive risk. Subsequent studies will assess cognition and use in-depth MRI to examine brain structure and function to more precisely align findings to specific VCID-relevant diseases.

Regional brain perfusion can be reduced in the brain for a number of reasons, including occlusive pathology, intrinsic cellular dysfunction, reduced neurovascular connectivity/function, as well as fewer vessels overall. TT male and CT and TT female mice show significantly reduced vascular density by immunohistochemistry at 6 months in the frontal cortex, as well as a trend in reduced density within the somatosensory and visual cortices at both 6 and 18 months of age. Because the phenotype appears by 6 months of age and does not appear to be age-related, this suggests a possible developmental difference/delay. It is feasible that aberrant signaling in the brain either disrupts initial vasculogenesis or encourages degeneration following development. GFAP-expressing astrocytes are significantly increased in male TT brains in both the visual and frontal cortices at 6 months, with a trend towards increased expression in 6 month old females and 18 month old males and females. Increased presence of activated astrocytes in regions where there is a noted reduction in vascular density could suggest the presence of an inflammatory microenvironment. This environment could contribute to or be a result of neurovascular dysfunction.

Evidence of vascular abnormalities as well as an increased astrocyte presence in the frontal cortex of CT and TT brains was further corroborated by electron microscopy. A significant number of imaged vessels (determined by morphology and presence of endothelial cells) demonstrated a severely reduced or absent lumen, which could contribute to reduced blood flow. Reduced luminal size may result from thickening of endothelial or mural cell layers, narrowing the diameter, or a dysfunction of these cells’ contractility. While some vascular cells could be experiencing intrinsic dysfunction that cannot be ascertained from ultrastructural imaging, others in CT and TT cortex demonstrated a clear apoptotic phenotype within endothelial cells and pericytes. Loss of endothelial cells and their tight junction proteins has long been associated with damage to the BBB. Cell death increases stress on surrounding endothelial cells to maintain the function of an intact barrier. Pericytes are also critical components of the neurovascular unit and play a key role in the regulation of blood flow, vessel contractility, and inhibition of vesicular transcytosis through endothelial cells. Pericyte dysfunction and dropout has been associated with Alzheimer’s disease in a number of studies^75–77^ with some demonstrating degeneration by observing pericyte debris in cerebral spinal fluid^75, 78^.

Astrocytes are known to support BBB integrity by maintaining the tight junctions between endothelial cells and supporting connections between neurons and vascular cells at the neurovascular unit^79, 80^. While they serve a protective role by containing damage, excessive or sustained astrocyte reactivity can contribute to chronic inflammation and neuronal dysfunction^81, 82^. It is unclear by EM whether the increase in astrocyte endfeet encircling vessels in the cortex of CT and TT mice represents a protective or damaging effect. The observed increase in microglia across brain regions also suggests an inflammatory environment, but again whether microglia are clearing debris from degenerating cells or inducing degeneration when interacting with the vascular cells remains to be determined.

In conclusion, the *Mthfr^677C>T^* mouse model demonstrates morphological and functional vascular deficits, enhanced glial cell activity and disrupted metabolism in the brain by 6 months of age. These findings represent a foundation from which to further explore phenotypes induced by the risk variant alone and also under different genetic and environmental conditions. A better understanding of lower-level genetic risk aids in building a more dimensional model to study the complex pathology of Alzheimer’s disease and other neurodegenerative diseases.

## Acknowledgements

We would like to thank Amanda Hewes and Melanie Maddocks for their day-to-day support, and Dylan Garceau, the MODEL-AD team, and JAX’s Genetic Engineering Technologies for development of the *Mthfr^677C>T^* model. We thank Todd Hoffert of JAX’s Clinical Assessment for establishing an in-house homocysteine assay. We would like to thank Paula Ashcraft and Brandi Wasek (Baylor Scott & White Research Institute, Dallas, Texas, USA), for the analysis of metabolites related to the folate and methionine cycles.

We would also like to acknowledge funding support from the Diana Davis Spencer Foundation (Howell), BrightFocus Foundation (grant no. A2020677F) (Reagan), the Canadian Institutes of Health Research (grant no. PJT173521) (Rozen), National Institute on Aging (grant no.U54AG054345) (MODEL-AD).

## Author contributions

AMR, LCG and GRH conceived the study. AMR established and aged the experimental cohorts, harvested tissue and performed immunohistochemistry, confocal imaging, electron microscopy. KEC and RR performed enzyme activity assays. AAB, KE, RS, LLF, SCP and PRT performed PET/CT. TB performed metabolite analysis. AMR and GRH wrote the manuscript that was edited and approved by all authors.

