## Supplemental Figure 1 for "The *677C>T* variant in *methylenetetrahydrofolate reductase* causes morphological and functional cerebrovascular deficits in mice"

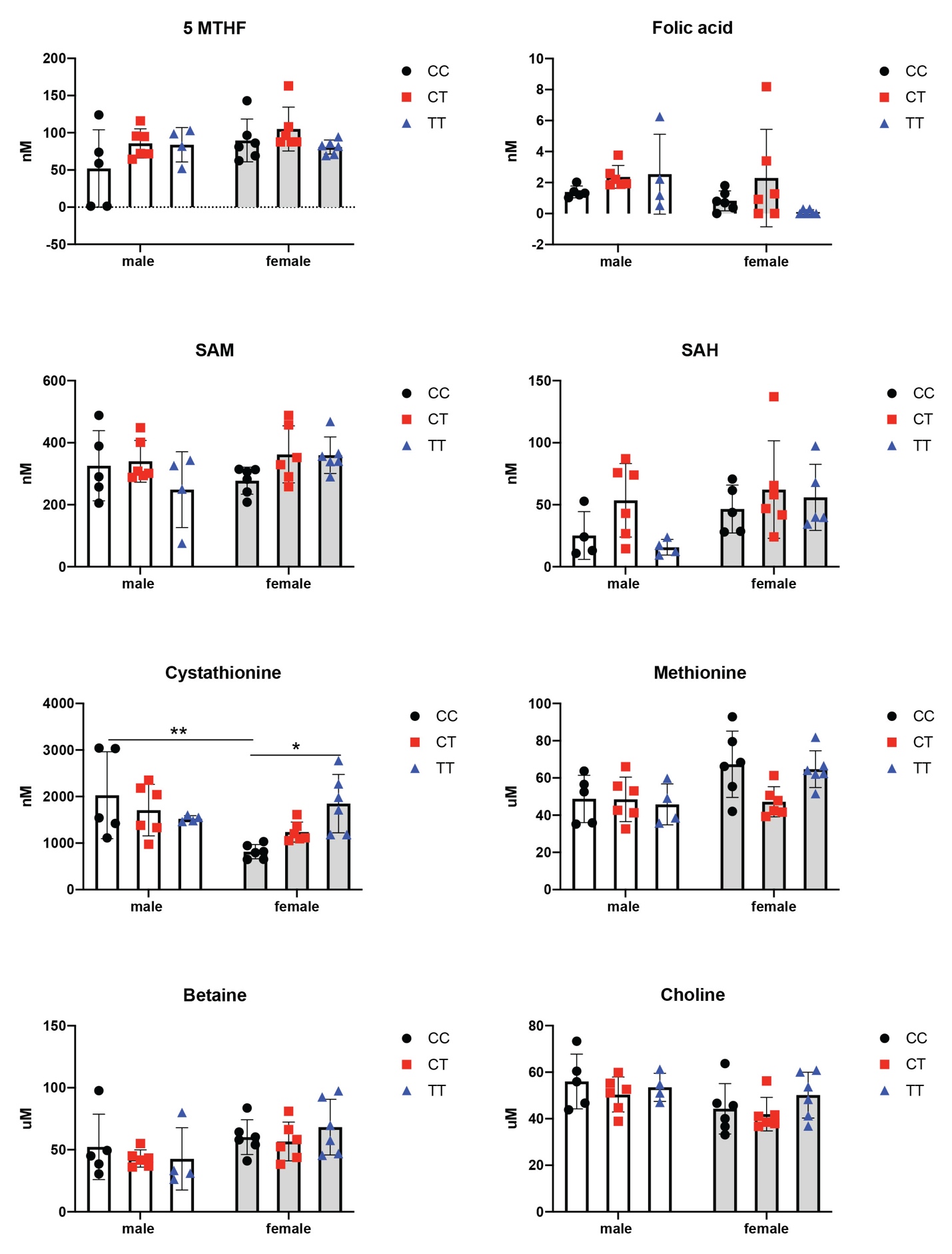

**Supplementary Fig 1.** Plasma from 3-4 month old male and female CC, CT and TT mice was analyzed for relevant metabolites in the folate and methionine cycles. Cystathionine is significantly increased in TT females compared to CC females. Male CC mice have significantly higher levels of cystathionine than CC females (P=0.0094 for interaction, P=0.0201 for sex; 95% CI {76.96-829.9}). No significant differences were observed in 5-MTHF, folic acid, SAM, SAH, methionine, betaine and choline (n=4-6/sex/genotype; Two-way ANOVA, Tukey post-hoc).

**Supplementary Table 1.** MANOVA results for all 27 brain regions analyzed by PET/CT (n=6/sex/genotype/age.) See excel file.
